# Ubiquitin-specific protease 20 promotes CCCP-induced mitophagy through deubiquitination and stabilization of serine/threonine protein kinase PINK1

**DOI:** 10.1101/2025.03.30.646148

**Authors:** Ga Hyun Park, Hye In Park, Donghyuk Shin, Kwang Chul Chung

**Author notes:** To whom correspondence should be addressed: Department of Systems Biology, College of Life Science and Biotechnology, Yonsei University, Yonsei-ro 50, Seodaemun-gu, Seoul 03722, Korea.

## Abstract

While Parkinson’s disease (PD) is predominantly sporadic, various mutations in the PTEN-induced putative kinase 1 (PINK1) gene have been linked to the autosomal recessive form of PD. PINK1, a serine/threonine protein kinase, holds a pivotal role in mitophagy - a process that selectively eliminates damaged mitochondria, overseeing mitochondrial quality control and ultimately safeguarding against neuronal cell loss in PD. Understanding the regulation of PINK1 stability is essential in comprehending PD pathology, given its involvement in a pro-survival pathway. Although some components of the ubiquitin-proteasome system (UPS) are recognized for mediating the proteolysis of PINK1, the specific enzyme(s) responsible for positively influencing PINK1 stability have remained elusive. In this study, we demonstrated that ubiquitin-specific protease 20 (USP20) functions as a novel deubiquitinating enzyme targeting PINK1. We found that USP20 positively regulates PINK1 levels by hydrolyzing Lys 48-linked polyubiquitin chains, promoting mitophagy under the treatment of mitochondrial depolarizing agent carbonyl cyanide *m*-chlorophenyl hydrazine (CCCP). Furthermore, CCCP treatment accelerates the deubiquitinating activity of USP20, facilitating the degradation of impaired mitochondria and enhancing mitochondrial quality control via PINK1 accumulation. Taken together, these findings unveil a novel enzyme, USP20, positively impacting PINK1 level and promoting CCCP-induced mitophagy. In addition, this study establishes a comprehensive map depicting how PINK1 can be regulated both positively and negatively through the coordinated action of multiple members in the UPS.

## Introduction

In Parkinson’s disease (PD), dopaminergic neurons primarily in a specific region of the midbrain called the substantia nigra are predominantly affected (Beitz, 2014). While various factors are implicated in causing PD, defective mitochondrial function and increased vulnerability to oxidative stress are considered its main pathogenic causes (Moon and Paek, 2015). Effective mitochondrial turnover through mitophagy is essential for maintaining neuronal health, and disruptions in this process are strongly associated with the development of PD (Liu et al., 2019). Although the majority of PD cases are sporadic, autosomal-dominant and recessive forms of familial PD, including mutations in α-synuclein, LRRK2, parkin, PTEN-induced kinase 1 (PINK1), DJ-1, and FBXO7, have also been identified (Shin and Chung, 2020; Park et al., 2018; Gao et al., 2022). Among these targets, PINK1 and parkin play crucial roles in mitophagy, a selective degradation of damaged mitochondria by autophagy and thus a critical response for the degradation of unhealthy mitochondria, and their ‘loss-of-function’ mutations result in mitochondrial dysfunction and defects in mitochondrial quality control, ultimately triggering neural degeneration in PD. Mitophagy is induced by various factors, including membrane depolarization, mitochondrial complex dysfunction, mutagenic stress, and proteotoxicity (Ge et al., 2020).

Upon damage to the mitochondrial membrane PINK1 accumulates on the outer mitochondrial membrane. Subsequently, PINK1 phosphorylates ubiquitin (Ub) and parkin, thereby stimulating the ubiquitin E3 ligase activity of parkin (Ge et al., 2020). Parkin then initiates a series of mitophagic reactions and eliminates defective mitochondria (Hu and Wang, 2016). Notably, the chemical mitochondrial uncoupler, carbonyl cyanide *m*-chlorophenyl hydrazine (CCCP), induces the loss of mitochondrial membrane potential (MMP), triggering PINK1-parkin-mediated mitophagy (Youle and Narendra, 2011; Ashrafi and Schwarz, 2013). Beyond these functions, PINK1 serves as a key enhancer of IL-1β signaling, facilitating the upregulation of inflammatory cytokines in response to IL-1β stimulation. Notably, our research demonstrates that PINK1 fosters interleukin-1β-mediated inflammatory signaling through positive regulation of both TRAF6-TAK1 and Tollip-IRAK1 interactions (Lee and Chung, 2012; Lee et al., 2012).

Despite the crucial functional and physiological roles of PINK1, the regulatory mechanisms influencing its stability remain largely unexplored. Currently, the ubiquitin-proteasome system (UPS) is acknowledged for impacting PINK1 stability. Notably, ubiquitin E3 ligases like UBR1, UBR2, and UBR4 participate in PINK1 degradation through the N-end rule pathway (Yamano and Youle, 2013). Additionally, we have identified chaperone-dependent CHIP as a novel E3 ubiquitin ligase of PINK1 (Yoo and Chung, 2017). Since ubiquitination generally operates reversibly, deubiquitinating enzymes (DUBs), including ubiquitin-specific proteases (USPs), play a crucial role in detaching ubiquitin from substrates (Meray and Lansbury, 2022; Li et al., 2022).

While multiple USPs are associated with diverse cellular mechanisms, their contribution to mitochondrial quality control remains elusive (Wang et al., 2022). For instance, USP33 deubiquitinates parkin, antagonizing PINK1-parkin-mediated mitophagy by deubiquitinating K63-linked poly-Ub chains (Niu et al., 2022; Durcan et al., 2014). In this context, maintaining PINK1 stability should also require a delicate balance between ubiquitinating and deubiquitinating enzymes. However, the reversed regulatory mechanism and the specific USPs involved are yet to be thoroughly investigated, guiding our ongoing efforts to identify the specific USP(s) biochemically and functionally associated with PINK1. Among the numerous USP members, USP20 interacts with PTEN and affects neuroinflammation via Toll-like receptor signaling pathway (Pan et al., 2022; Jean-Charles et al., 2016). For example, USP20 promotes IL-1β-evoked signaling in vascular inflammation through IRAK1-mediated phosphorylation (Zhang et al., 2023). Furthermore, USP20 targets TRAF6 to negatively regulate NF-κB signaling (Yasunaga et al., 2011).

Based on the previous findings that indicate the common involvement of USP20 and PINK1 in both PTEN- and Toll-like receptor-signaling pathways, this study aimed to investigate the potential linkage and mechanisms connecting these two proteins. Consequently, we demonstrated that USP20 functions as a novel deubiquitinating enzyme targeting PINK1, promoting its accumulation by hydrolyzing K48-linked poly-Ub chains. Through the prevention of PINK1 degradation via UPS, USP20 has the potential to enhance mitophagy in response to CCCP exposure. Since there has been no direct association established between USP20 and mitochondrial dynamics, our investigation delves deeper into exploring the specific connection between USP20 and PD.

## Materials and Methods

### Cell culture and DNA transfection

Human embryonic kidney 293 (HEK293) cells were maintained in DMEM supplemented with 10% fetal bovine serum and 100 U/mL penicillin-streptomycin. Mouse embryonic fibroblasts (MEFs) and human neuroblastoma SH-SY5Y cells were maintained in DMEM/F12 supplemented with 10% FBS and 100 U/mL penicillin-streptomycin. The cells were grown at 37°C in 5% CO_2._ All DNA transfections were performed by using Lipofectamine 2000 or Mirus reagent, according to the manufacturer’s instructions. The PINK1-knockout and control MEFs were kindly provided by J. Shen (Harvard Medical School, Boston, MA, USA).

### DNA constructs and RNA interference

The plasmids encoding Myc-tagged human wild-type PINK1 (pBOs-3x-Myc-hPINK1-WT) and Flag/HA-tagged USP20 (pDEST-Flag/HA-USP20) were kindly provided by J. Chung (Seoul National University, Seoul, Korea) and J. Song (Yonsei University, Seoul, Korea), respectively. The plasmid encoding GST-tagged PINK1 kinase domain (amino acids 112–496; pGEX5T-1-PINK1_112-496_) was provided by M.R. Cookson (National Institutes of Health). The plasmids encoding Flag-tagged deletion mutants of USP20 (USP20_1-144_, USP20_145-647_, and USP20_648-895_) were a kind gift from S.H. Park (Sungkyunkwan University, Suwon, Gyeonggi-do, Korea). The plasmids encoding Myc-tagged deletion mutants of PINK1 (PINK1_111-581_, PINK1_156-581_, and PINK1_300-581_) were constructed as previously described (14). To construct the plasmid encoding catalytic inactive USP20 mutant (USP20-C154S), site-directed mutagenesis was performed using the QuickChange XL-site directed mutagenesis kit (Stratagene) and pDEST-Flag/HA-USP20 as a template. All plasmid sequences were confirmed by using DNA sequencing (Bionics, Seoul, Korea). Small interfering RNAs (siRNAs) for USP20 and control scrambled siRNA (cat. #51-01-14-04) were purchased from Integrated DNA Technologies (Hanam-si, Gyeonggi-do, Korea). The USP20-specific siRNA duplex sequences were as follows: 5′-CGACACCUUCAUCAAGUUGAACAAG-3′ (sense) and 5′-UAGCUGUGGAAGUAGUUCAACUUGUUC-3′ (anti-sense).

### Co-immunoprecipitation (co-IP) and immunoblot analysis

Cells were scraped and washed extensively with ice-cold phosphate-buffered saline (PBS). Cells were then mixed with 1% of Nonidet P-40 lysis buffer (50 mM Tris, pH 7.5, 1% Nonidet P-40, 150 mM NaCl, and 10% glycerol) including the protease inhibitor cocktail (1 mM NaF, 1 μg/mL leupeptin, 1 μg/mL aprotinin, 1 mM Na_3_VO_4_, and 0.2 mM PMSF), incubated in the ice for 10 min, and centrifuged at 13,000 × *g* for 15 min at 4°C. For immunoprecipitation, 500∼1000 μg of cell lysates were incubated with 1 μg of appropriate antibodies overnight at 4°C with gentle rotation. Protein A-Sepharose beads were then incubated for 2 h at 4°C and washed three times with lysis buffer. The immunocomplexes were eluted in 2X sample buffer and dissociated by boiling for 5 min. The samples were separated by SDS-PAGE gels and transferred to a nitrocellulose membrane (Whatman, GE Healthcare Life Sciences). Membranes were blocked for 1 h at room temperature in TBST buffer (20 mM Tris, pH 7.5, 137 mM NaCl, and 0.1% Tween 20) with 5% nonfat dry milk, and incubated with the appropriate primary antibody at 4°C overnight. Membranes were washed with TBST, incubated in HRP-conjugated secondary IgG antibody for 2 h room temperature. The protein bands were detected by using ECL reagents (Abclon), following the manufacturer’s instructions.

### Immunocytochemistry analysis

HEK293 cells were seeded onto poly-L-lysine-coated cover glasses in 6-well plates. After DNA transfection, cells were washed three times with PBS (pH 7.4) and fixed with 3.7% formaldehyde for 10 min. After fixation, cells were permeabilized with 0.1% Triton X-100 for 10 min, blocked with 1% BSA in TBST for 1 h at room temperature, and immunostained using rabbit polyclonal anti-PINK1 and/or rabbit polyclonal anti-USP20 antibodies. The samples were then incubated with Alexa Fluor 488- or Alexa Fluor 594-conjugated anti-IgG antibodies. Microscopic images were obtained using the LSM 880 confocal microscope (Carl Zeiss, Oberkochen, Germany) and processed using the Zeiss LSM Image Browser (Carl Zeiss).

### Alphafold prediction of PINK1:USP20:Ubiquitin complex

To obtain molecular details of the interaction between PINK1 and USP20, we performed AlphaFold3-server-based PINK1:USP20 complex prediction (alphafoldserver.com/about) (Abramson et al.,2024). The kinase domain of PINK1 (PINK1_115-581_), the USP catalytic domain (USP20_145-647_), and full-length ubiquitin sequences were used for prediction. To avoid any steric hindrance caused by the disordered region of USP20 (USP20_250-441_), we decided to use the core catalytic domain of USP20 (USP20_145-647_-Δ250-441). The AlphaFold3 server generated five complex structures with high similarity, and the representative model was further analyzed using PyMOL (The PyMOL Molecular Graphics System, Version 3.0 Schrödinger, LLC).

### Proximity Ligation Assay

A PLA was performed utilizing the Duolink PLA assay kit (DUO92101, Sigma-Aldrich) according to the manufacturer’s protocol. The PLA signals are shown in red, while the nuclei are indicated in blue.

### In vitro GST pull-down assay

Bacterial plasmids encoding GST-PINK1-112-496 (pGEX5T-1-PINK1-112-496) or GST (pGEX4T-1-GST) were transformed into *E. coli* BL21 cells (Thermo Fisher Scientific). Recombinant GST proteins was induced by the addition of 0.1 mM of isopropyl 1-thio-β-D-galactopyranoside (IPTG) for 16 h at 16°C. Bacterial pellets were lysed with lysis buffer and sonicated until transparent. The resulting supernatant was incubated with Glutathione Sepharose 4D for 16 h at 4°C. Beads were then washed with PBS buffer and proteins were eluted with elution buffer. For USP20 preparation, HEK293 cells were transiently transfected for 24 h with Flag-USP20, harvested, and lysed in lysis buffer. Cell lysates containing Flag-USP20 protein were incubated with immobilized GST or GST-tagged PINK1 proteins overnight at 4°C. After washing, samples were analyzed by western blotting after addition of 2X sample buffer.

### Analysis of mitochondrial membrane potential

The mitochondrial membrane potential was analyzed using JC-1 dyes (Life Technologies) according to the manufacturer’s instructions. HEK293 cells were placed stained with 0.5 μg/ml JC-1 dyes (Invitrogen, MA, USA) for 25 min at 37°C in 5% CO_2_. Cells were washed three times with PBS (pH 7.4) and analyzed using the LSM 800 confocal microscope.

### Statistical Analysis

All statistical analyses were performed using an unpaired student’s *t*-tests and one-way ANOVA. All values are reported as mean ± S.D. of at last three independent experiments. The GelQuant.NET software (version 1.8.2; biochemlabsolutions.com) was used for measuring the Western blot gel intensities.

## Results

### PINK1 interacts with USP20 in mammalian cells

Building on earlier findings that indicated the shared involvement of USP20 and PINK1 in both PTEN- and Toll-like receptor-signaling pathways, we aimed to investigate whether USP20 could function as a novel regulator of PINK1. If so, we further assessed the physiological consequences of USP20 following PINK1 modification. To test this hypothesis, we initially examined the binding interaction between these two proteins in mammalian cells. HEK293 cells were transfected with the plasmid encoding Myc-tagged PINK1 alone or in combination with Flag/HA-tagged USP20. Cell lysates were immunoprecipitated with anti-Myc antiserum, and immunoblotting with anti-Flag antibodies revealed that ectopically expressed USP20 binds to PINK1 (Figure 1A). Repeating the co-immunoprecipitation (co-IP) assay in reverse yielded the same result (Figure 1B). Subsequently, we performed the co-IP assay with their respective endogenous antiserum, confirming the interaction between these two proteins in both HEK293 and SH-SY5Y cells (Figure 1C, D). Finally, immunostaining of cells demonstrated the co-localization of these two proteins in the cytoplasm (Figure 1E). Next, we confirmed the direct interaction between these two proteins by performing an *in vitro* GST pull-down assay (Figure 1F). To validate the endogenous interaction, we additionally conducted a Proximity Ligation Assay (PLA), which demonstrated that endogenous PINK1 interacts with USP20 in the cytosol (Figure 1G). Overall, our data indicate that USP20 binds to PINK1 in mammalian cells.

**Figure 1.**
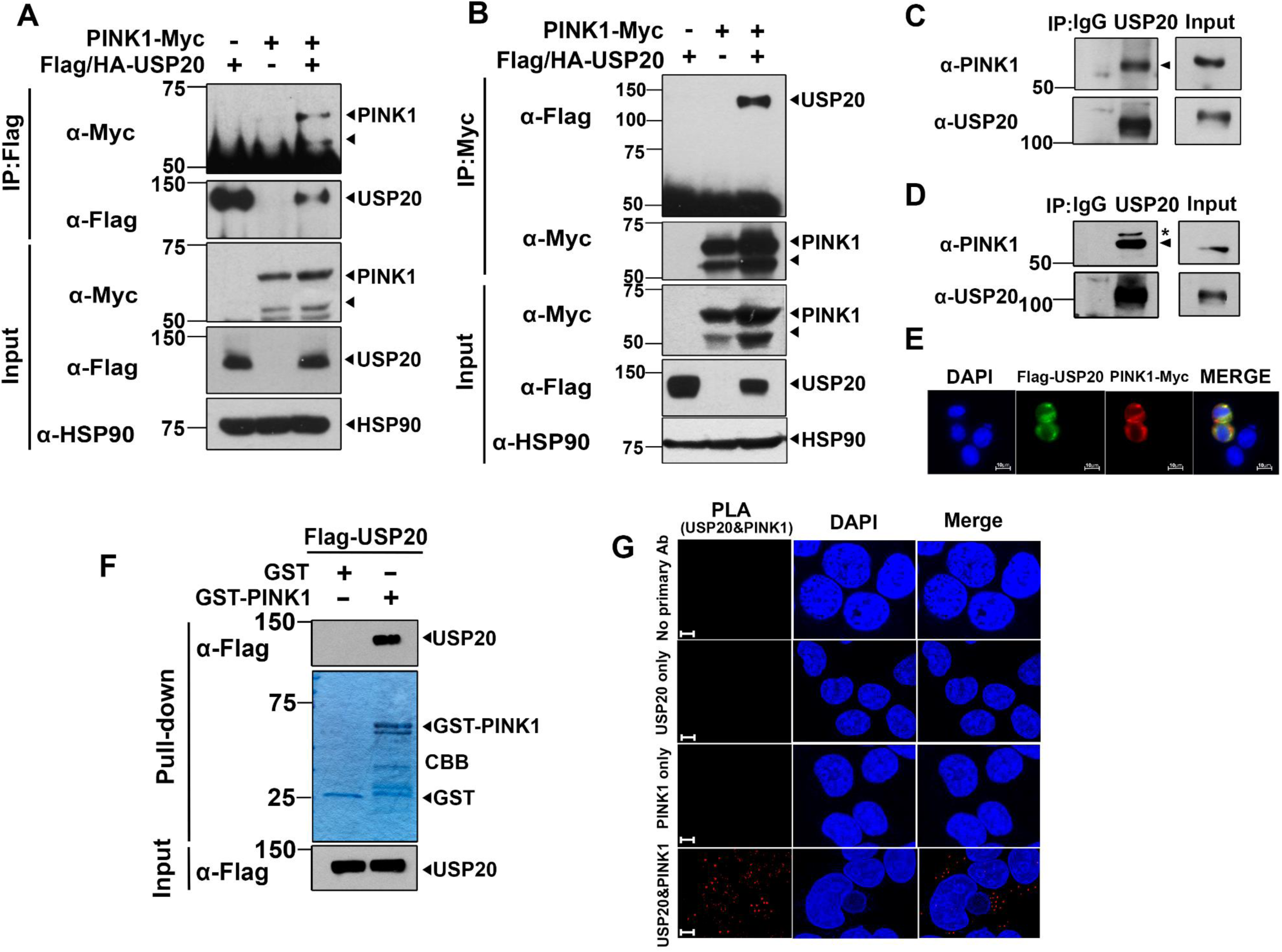
USP20 interacts with PINK1. **(A)** HEK293 cells were transfected for 24 h with a plasmid encoding Myc-tagged PINK1 or/and Flag/HA-USP20. Cell lysates were immunoprecipitated with an anti-Flag antibody, followed by immunoblotting with the indicated antibodies. **(B)** Cell lysates were immunoprecipitated with an anti-Myc antibody, followed by immunoblotting with the indicated antibodies. HSP90 served as a loading control. **(C)** HEK293 cell lysates were immunoprecipitated using either IgG (control) or anti-USP20 antibody, followed by immunoblotting with the indicated antibodies. **(D)** SH-SY5Y cell lysates were immunoprecipitated using either anti-USP20 antibody or IgG, followed by immunoblotting with the indicated antibodies. Asterisks indicate non-specific bands. **(E)** HEK293 cells were transfected with plasmids expressing Flag-USP20 and/or Myc-PINK1. The transfected cells were stained using anti-Flag (Green) and anti-Myc antibody (Red). Nuclei were counterstained with DAPI (blue). Scale bars = 10 μm. **(F)** The *in vitro* GST pull-down assay was performed using purified GST or GST-PINK1. Reaction mixtures were added to cell lysates derived from HEK293 cells transfected with Flag-USP20. After washing the beads, they were analyzed by immunoblotting using anti-Flag antibody. The purified GST or GST-PINK1 proteins were stained with Coomassie brilliant blue (CBB) and shown in the middle panel. **(G)** The PLAs were conducted using the primary antibodies targeting PINK1 and USP20. The red dot indicates the interaction between endogenous PINK1 and USP20. Scale bars = 10 μm.

### PINK1 kinase domain interacts with the catalytic domain of USP20

Next, we wondered how PINK1 is recognized by USP20. PINK1 is composed by 581 amino acids (a.a.), encompassing the mitochondrial targeting sequence (MTS), transmembrane region (TM), N-terminal regulatory region (NT), N-lobe of the kinase domain (N-lobe), C-lobe of the kinase domain (C-lobe), and the C-terminal domain (CTD). USP20, with a length of 847 a.a., contains four distinct structural domains, which include the N-terminal zinc-finger ubiquitin binding domain (Zf-UBP), the catalytic domain (USP), and two tandem DUSP domains (DUSP1 and DUSP2). We performed AlphaFold3-based complex structure prediction (Abramson et al., 2024). The predicted complex structure of ubiquitin, the kinase domain of PINK1 (PINK1_115-581_), and the catalytic core domain of USP20 encompassing a.a. 145-647 with a deletion of a.a. 250-441 (USP20_145-647_-Δ250-441) indicates that both the N-lobe and C-lobe of the PINK1 kinase domain make contacts with USP20 (Figure 2A).

**Figure 2.**
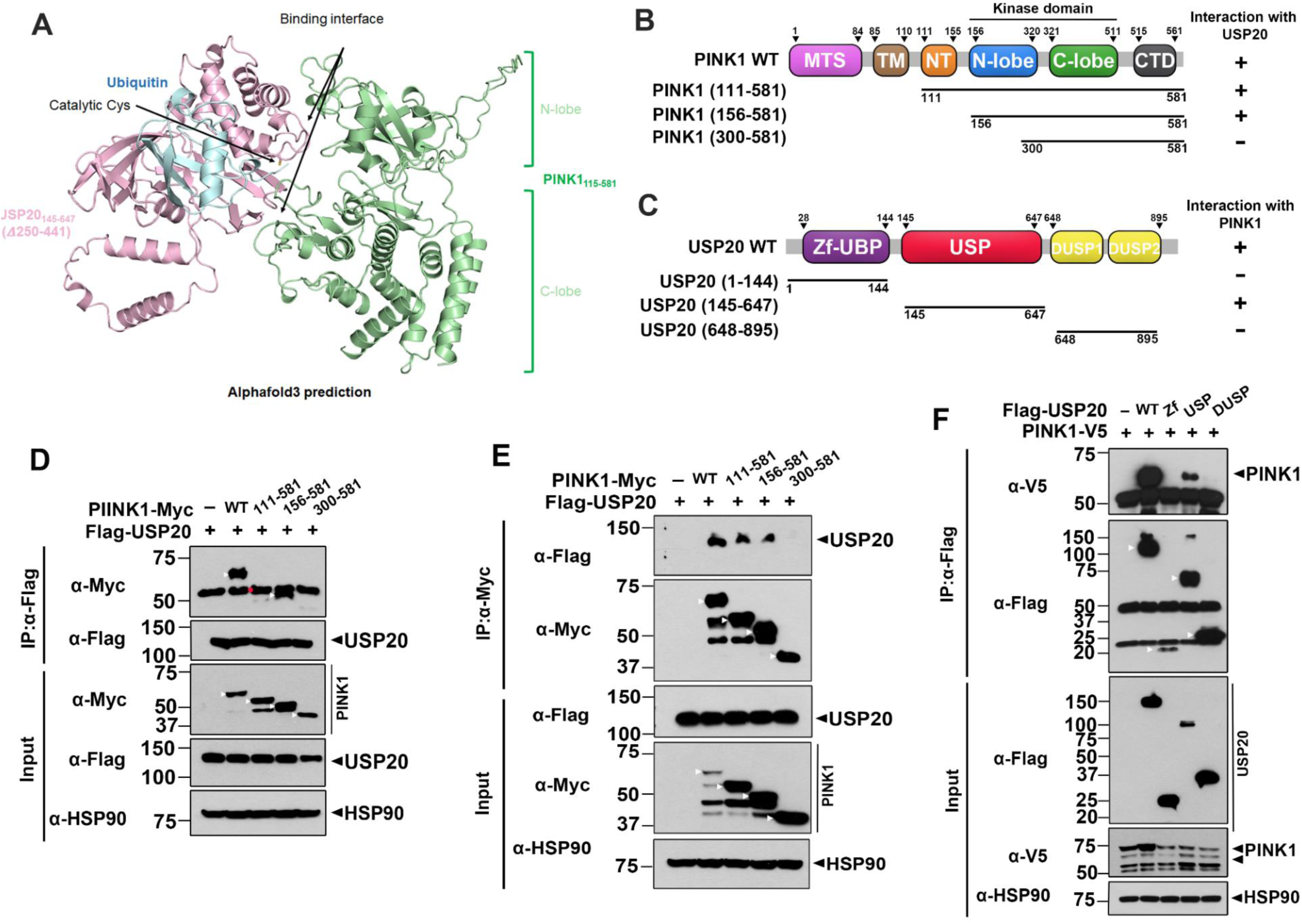
The intact kinase domain of PINK1 and the catalytic domain of USP20 are important for binding. **(A)** The representative model of AlphaFold3-predicted complex structure of ubiquitin (cyan), the kinase domain of PINK1 (PINK1_115-581,_ pale green), and the catalytic core domain of USP20 (USP20_145-647-_Δ250-441, pink). Catalytic cysteine of USP20, N-lobe, and C-lobe of PINK1 are depicted. (B and C) Schematics of full-length PINK1 (PINK1-WT) and its deletion mutants **(B)**, as well as full-length USP20 (USP20-WT) and its deletion mutants **(C)**, are shown. The results from the co-immunoprecipitation and binding assays between the full-length or deletion mutants of USP20 and PINK1 are displayed on the right. The (+) indicates binding, while (-) indicates no binding. **(D and E)** Where indicated, HEK293 cells were transfected for 24 h with plasmids encoding Flag-tagged wild-type USP20, Myc-tagged full-length PINK1, or one of its deletion mutants alone or in combination. Cell lysates were immunoprecipitated using either anti-Flag **(D)** or anti-Myc **(E)** antibodies, followed by immunoblotting with the indicated antibodies. (F) Where indicated, HEK293 cells were transfected for 24 h with plasmids encoding V5-tagged wild-type PINK1, Flag-tagged full-length USP20, or one of its deletion mutants alone or in combination. Cell lysates were immunoprecipitated using anti-Flag antibody, followed by immunoblotting with the indicated antibodies. HSP90 served as a loading control.

To validate this structural prediction and identify the specific domains of USP20 and PINK1 responsible for binding, we generated several deletion mutants, each lacking conserved functional and/or structural domains of the respective protein (Figure 2B, C). We examined the predicted binding interactions using PINK1 constructs containing the intact kinase domain and NT, and CTD (PINK1_111-581_), the kinase domain with CTD (PINK1_156-581_), and the C-lobe of the kinase domain with CTD (PINK1_300-581_) to assess their binding to Flag-USP20. PINK1 interacts with USP20 when the intact kinase domain of PINK1 is present (Figure 2D). However, deletion of the N-lobe of the kinase domain abolished the PINK1 binding to USP20. The predicted complex structure is further supported by examining the interaction of different domains on USP20 with PINK1 (Figure 2E). Consistent with the predicted structure, the catalytic USP domain interacts with PINK1, while the Zf-UBP or DUSP1/2 domains did not (Figure 2F). These findings suggest that the intact kinase domain of PINK1 is required for its binding to the catalytic domain of USP20.

### USP20 causes the increase of PINK1 level

USP20 operates as an ubiquitin protease, exerting control over the ubiquitination process and subsequent proteasome-mediated degradation of its target substrates. To gain further insight as to how USP20 and PINK1 are linked, we examined how USP20 influences the protein stability of PINK1. Upon transfecting HEK293 cells with PINK1-Myc or increasing doses of USP20 alone or in combination, our immunoblotting analysis revealed a dose-dependent increase in both endogenous and exogenous PINK1 levels with the overexpression of USP20 (Figure 3A, B). This consistent result was further confirmed in other mammalian cells, such as SH-SY5Y and A549 cells, where the presence of USP20 led to an increase in endogenous PINK1 levels (Supplementary Figure S1A).

**Figure 3.**
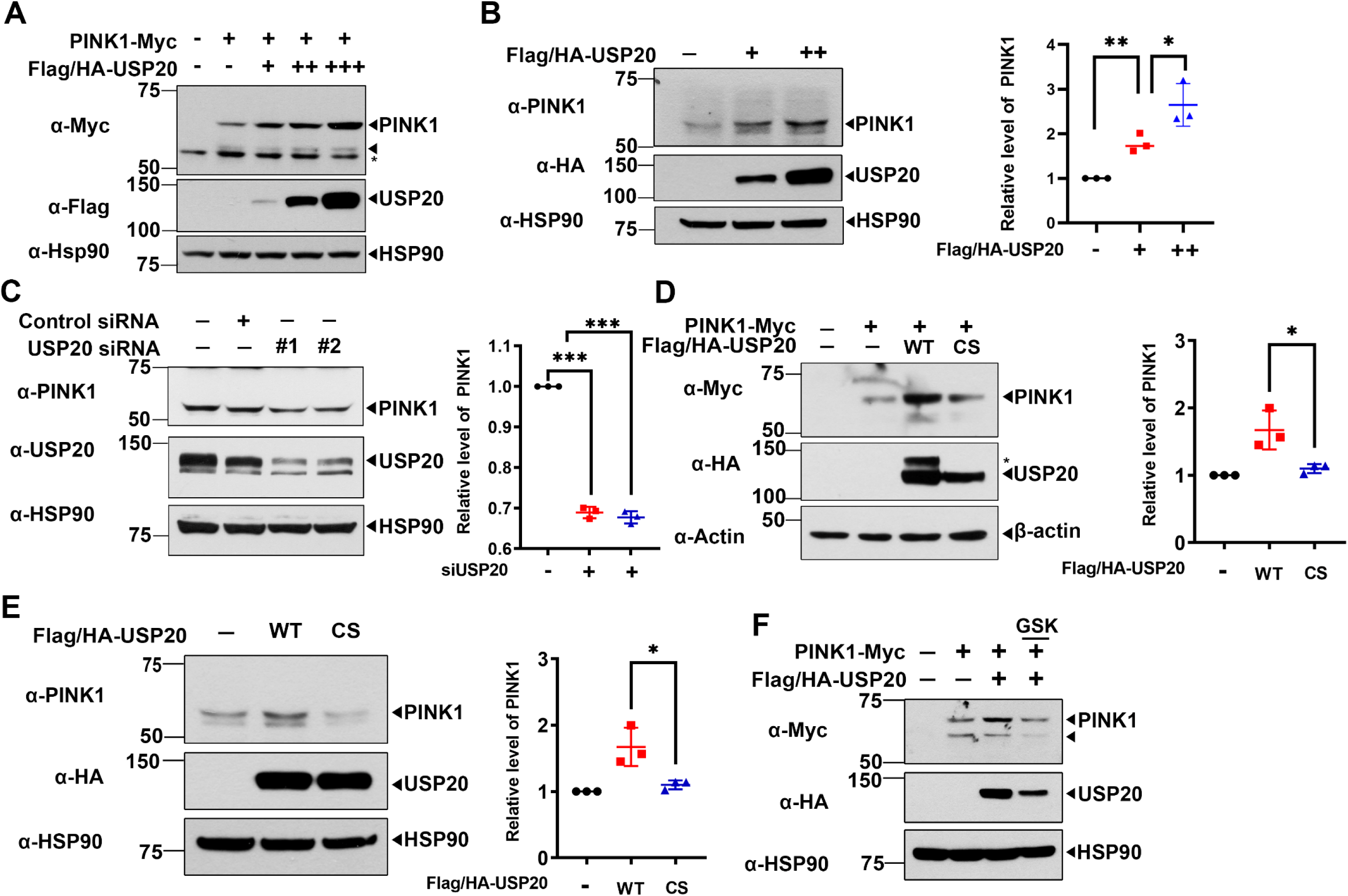
USP20 increases the protein levels of PINK1. **(A)** HEK293 cells were transfected for 24 h with the plasmids encoding wild type PINK1-Myc alone or together with increasing amounts of a plasmid encoding Flag/HA-USP20, and cell lysates were immunoblotted with anti-Myc or anti-HA antibodies. Asterisks indicate non-specific bands. **(B)** HEK293 cells were transfected for 24 h with increasing amounts of a plasmid encoding Flag/HA-USP20. Cell lysates were immunoblotted with anti-PINK1 or anti-HA antibodies. Relative levels of PINK1 were quantified, and the results are presented as the mean ± S.D. of three independent experiments (\*\**p* ≤ 0.001; \**p* ≤ 0.05). **(C)** HEK293 cells were transfected for 48 h with control-siRNA or *USP20*-siRNA. Cell lysates were immunoblotted with anti-PINK1 or anti-USP20 antibodies. Relative PINK1 levels were quantified and the results are presented as the mean ± S.D. of three independent experiments (\*\*\**p* ≤ 0.0001). **(D)** HEK293 cells were transfected for 24 h with a plasmid encoding PINK1-Myc, Flag/HA-USP20-WT, Flag/HA-USP20-C154S (CS) alone or in combination. Relative PINK1 levels were quantified and the results are presented as the mean ± S.D. of three independent experiments (\*\*\**p* ≤ 0.0001; \**p* ≤ 0.05). Asterisks indicate non-specific bands. **(E)** HEK293 cells were transfected with plasmid encoding Flag/HA-USP20-WT or Flag/HA-USP20-C154S (CS). Cell lysates were immunoblotted with anti-PINK1 or anti-HA antibody. Relative PINK1 levels were quantified, and the results are presented as the mean ± SD of three independent experiments (\**p* ≤ 0.05). **(F)** HEK293 cells were transfected for 24 h with plasmids encoding wild type PINK-Myc and/or Flag/HA-USP20-WT. Cells were treated for 24 h with 10 μg/ml GSK2643943A, and cell lysates were immunoblotted with the indicated antibodies. HSP90 and actin served as loading control.

Furthermore, we explored the impact of siRNA-mediated knockdown of *USP20* on PINK1-Myc levels, confirming the efficiency of USP20-knockdown through immunoblotting analysis. In contrast to the effect of USP20 overexpression, suppressing USP20 resulted in a reduction of PINK1-Myc levels (Figure 3C). Intriguingly, this effect was nullified in the presence of a catalytic-inactive USP20 mutant with a substitution of Cys-154 with serine, demonstrating no discernible impact on exogenous and endogenous PINK1 levels (Figure 3D, E). Next, we explored the impact of the chemical inhibitor GSK2643943A, specifically targeting USP20, on PINK1-Myc levels (Figure 3F). As anticipated, pretreatment of cells with 10 μM of GSK2643943A for 24 h obstructed the positive modulatory effect of USP20 on PINK1-Myc levels, leaving the basal level of PINK1 unchanged (Figure 3F).

In summary, these findings suggest that USP20 plays a pivotal role in promoting PINK1 levels with the catalytic activity of USP20 proving critical for the accumulation of PINK1.

### USP20 positively regulates the protein stability of PINK1

To elucidate the impact of USP20 on modulating the protein stability of PINK1, we conducted a series of experiments involving HEK293 cells. Following a 24-hour transfection with plasmids encoding Flag/HA-tagged USP20 or its catalytically inactive mutant (Flag/HA-USP20-CS), either individually or in conjunction with PINK1-Myc, cells were subjected to a time-dependent treatment with 100 μM cycloheximide. Co-expression of wild-type PINK1 and wild-type USP20 revealed a noteworthy extension in the half-life of PINK1, which is attributed to USP20 (Supplementary Figure S2A). However, no such effect was observed with the catalytically inactive mutant of USP20 (Supplementary Figure S2A). Conversely, immunoblot analysis demonstrated a reduction in the half-life of endogenous PINK1 upon siRNA-mediated *USP20*-knockdown (Supplementary Figure S2B). These findings collectively underscore the significant role of USP20 in intricately and positively regulating the protein stability of PINK1.

### USP20 deubiquitinates PINK1 via targeting K48-linked polyubiquitin chains

Given the substantial increase in PINK1 protein levels resulting from the deubiquitinating activity of USP20, we subsequently investigated whether USP20 directly deubiquitinates PINK1. Given that K48-linked and K63-linked Ub chains are primarily associated with proteasome degradation and various signaling pathways, respectively, we focused on these two Ub linkages to identify the specific effect of USP20 on PINK1 polyubiquitination (Zhang et al., 2013). To confirm this, HEK293 cells were transfected for 24 h with a plasmid encoding PINK1-Myc, either alone or together with a plasmid encoding USP20-WT or its catalytically inactive USP20-CS mutant, and an immunoprecipitation assay was performed (Figure 4A). As expected, the wild-type USP20 significantly decreased the polyubiquitination of PINK1, whereas the USP20-CS mutant did not (Figure 4A). Additionally, we investigated which polyubiquitination chain of PINK1 was deubiquitinated by USP20. HEK293 cells were transfected with wild-type USP20, and anti-K48 or anti-K63 antibodies were used to detect the poly-Ub-linked lysine residues. The immunoblot results revealed that the K48-ubiquitin linkage regulates the deubiquitination of PINK1, but not via the K63-ubiquitin chain (Figure 4B, C). These results indicate that USP20 directly mediates the polyubiquitination of PINK1 by targeting K48-linked polyubiquitin chains.

**Figure 4.**
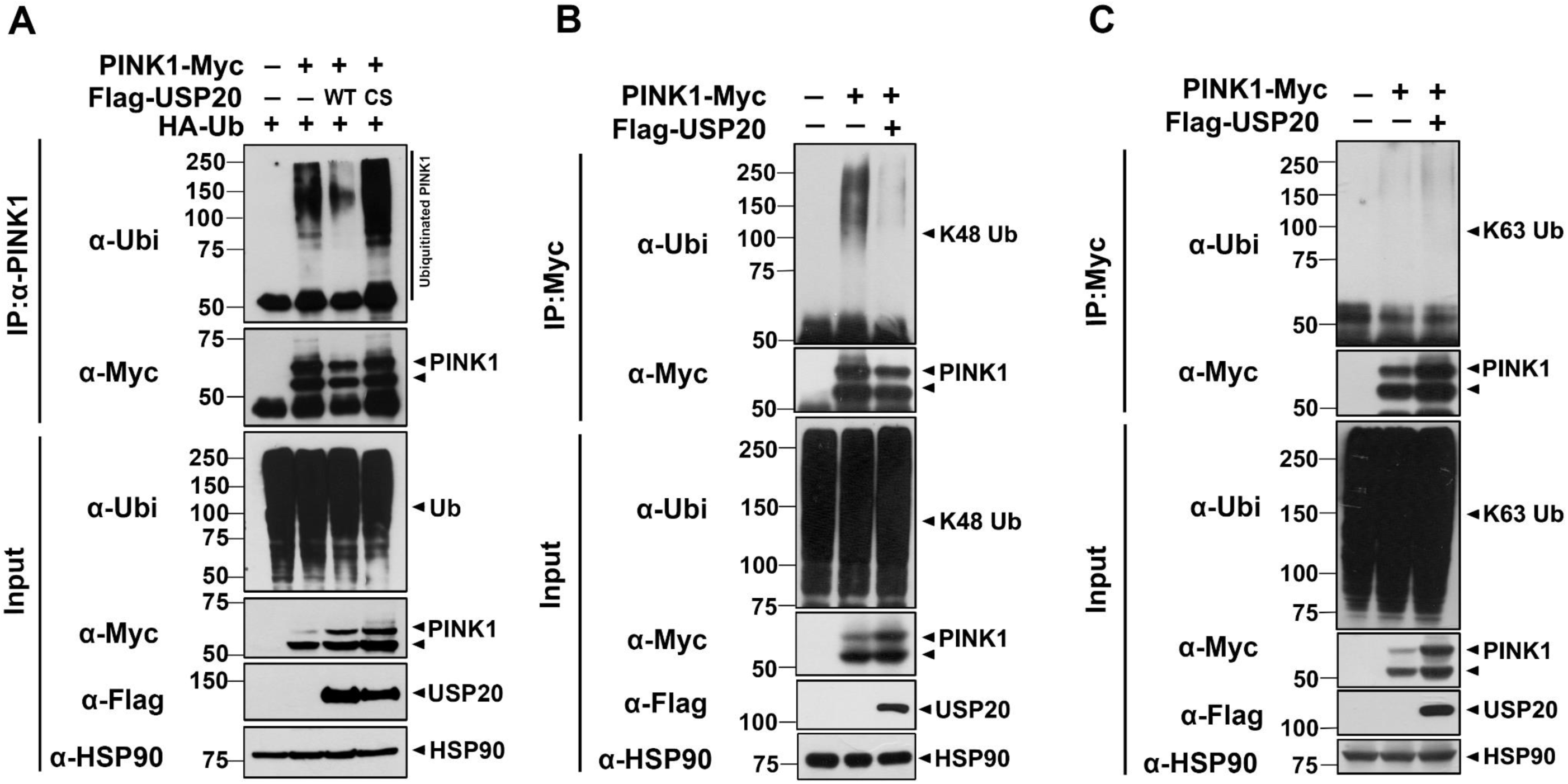
USP20 deubiquitinates PINK1 by targeting K48-linked poly-Ub chain. **(A)** Where indicated, HEK293 cells were transfected for 24 h with plasmids encoding PINK1-Myc alone or in combination with Flag-USP20-WT or Flag-USP20-CS, and treated for an additional 6 h with 10 μM of MG132. Cell lysates were immunoprecipitated using anti-PINK1 antibody, followed by immunoblotting with the indicated antibodies. **(B, C)** Where specified, HEK293 cells were transfected for 24 h with plasmids with encoding PINK1-Myc alone or in combination with Flag-USP20, and treated for an additional 6 h with 10 μM MG132. Cell lysates were immunoprecipitated with anti-K48 or anti-K63 antibodies, and the precipitates were immunoblotted with the indicated antibodies. HSP90 served as a loading control.

### USP20 enhances autophagy marker levels via PINK1 accumulation under CCCP treatment

Since mitophagy is a selective process involving the mitochondrial degradation via PINK1 recruitment and autophagic flux, we further examined how USP20 affects autophagy marker levels under treatment of mitochondrial uncoupler CCCP. CCCP is commonly used to reduce mitochondrial membrane potential, leading to the loss of mitochondria membrane integrity and its depolarization, which mainly contributes to the progression of PD. HEK293 cells were transfected with a plasmid encoding the catalytically inactive USP20-CS mutant and/or treated with CCCP. Immunoblot analysis of the lysates showed that the LC3-II level increased under CCCP treatment and slightly decreased with the overexpression of the USP20-CS mutant (Figure 5A). Additionally, p62/SQSTM1, known as an autophagic protein co-localized with depolarized mitochondria, leading to accumulation in the cytosol and subsequent recruitment of autophagic machinery (Ivankovic et al., 2016; Tsefou and Ketteler, 2022). Similar to the previous effect, the endogenous p62 level also exhibited an increase with CCCP treatment, and it was further elevated with USP20-WT (Figure 5B). Immunocytochemical analysis revealed that the LC3 level increased in the presence of USP20 compared to mock-transfected cells (Figure 5C).

**Figure 5.**
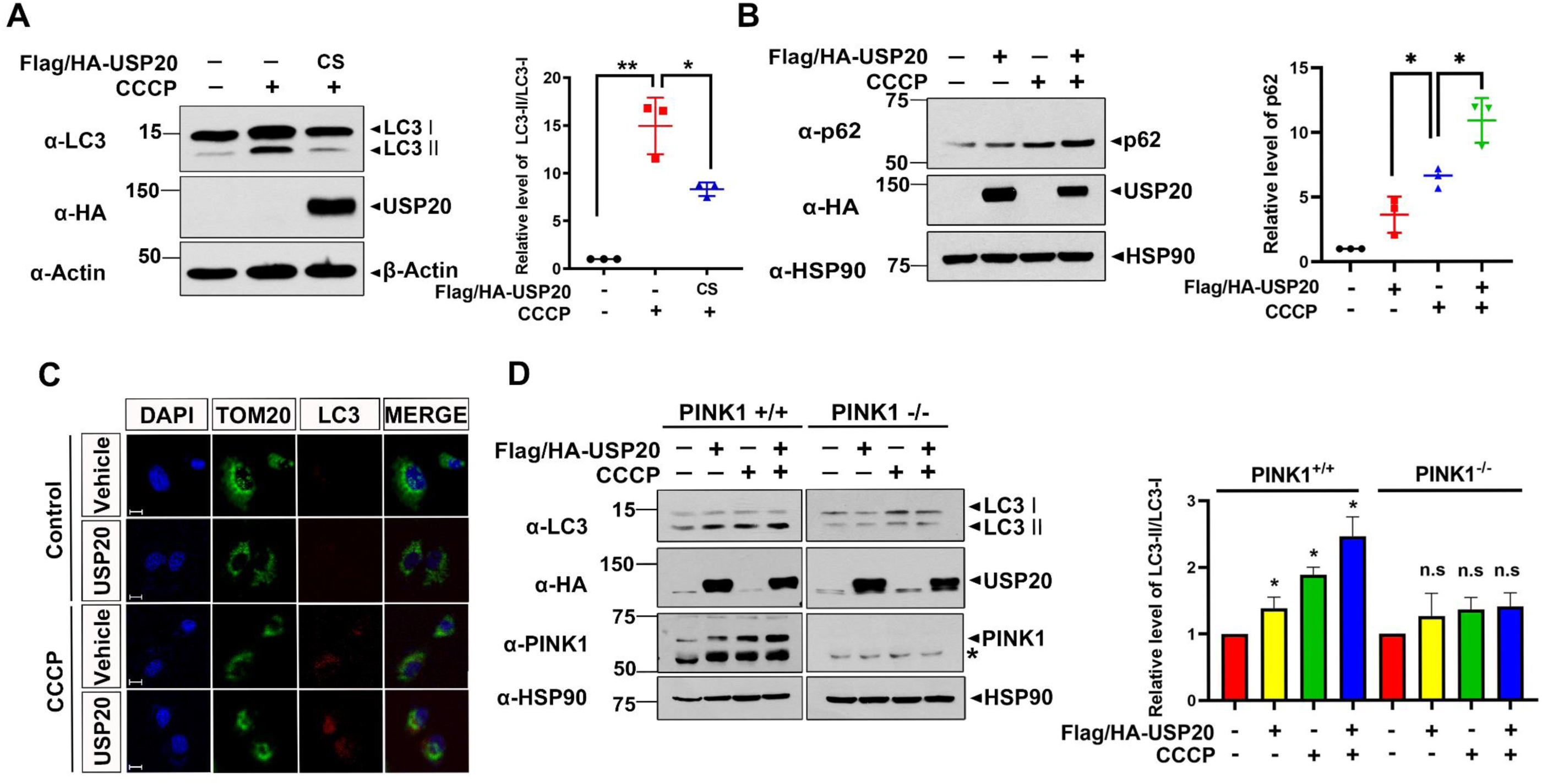
Autophagy marker levels are increased by USP20 under CCCP treatment. **(A, B)** HEK293 cells were transfected for 24 h with a plasmid encoding Flag/HA-USP20-WT or Flag/HA-USP20-CS, and treated with 10 μM CCCP for 4 h. Cell lysates were immunoblotted with the indicated antibodies. Relative levels of LC3 and p62 were quantified and the results are presented as the mean ± S.D. of three independent experiments (\*\**p* ≤ 0.001; \**p* ≤ 0.05). **(C)** HEK293 cells were mock-transfected or transfected with plasmid encoding Flag/HA-USP20 for 24 h, and treated with 10 μM CCCP for 6 h. Cells were then stained and fixed with anti-Tom20 (Green) or anti-LC3 (Red) antibodies, and DAPI for the nuclei. **(D)** Where indicated, PINK1^+/+^ and PINK1^-/-^ MEFs were transfected for 24 h with a plasmid encoding Flag/HA-USP20, and treated with 10 μM CCCP for 4 h. Relative levels of LC3 were quantified and the results are presented as the mean ± S.D. of three independent experiments (\*\**p* ≤ 0.001; \**p* ≤ 0.05). Asterisks indicate non-specific bands. Actin and HSP90 served as loading controls.

Next, we investigated whether the presence of PINK1 was essential for the USP20-mediated increase in these autophagy markers. In PINK1^+/+^ MEF cells, the LC3-II level gradually increased with CCCP treatment, but PINK1^-/-^ cells showed no significant change in LC3-II levels (Figure 5D). These results suggest that USP20-mediated accumulation of PINK1 is essential for the increase in autophagy markers. Taken together, the data suggest that USP20 increases autophagy marker levels by accumulating PINK1 levels.

### USP20 mediates the deubiquitination of PINK1 after treatment with CCCP

We then examined whether USP20 could elevate PINK1 levels upon exposure to CCCP through PINK1 deubiquitination. After HEK293 cells were treated with vehicle or CCCP, immunoblot analysis of cell lysates revealed that the endogenous USP20 level was unaffected by CCCP treatment (Figure 6A). We subsequently explored whether PINK1 directly binds to USP20 in the absence or presence of CCCP treatment. Co-IP analysis revealed that ectopically expressed PINK1 interacts with USP20 in the basal state. Moreover, CCCP treatment had no further effect on the binding affinity between PINK1 and USP20 (Figure 6B). Building on our previous finding that USP20 significantly reduces PINK1 polyubiquitination, we investigated whether the deubiquitinating activity of USP20 toward PINK1 is increased by CCCP treatment. As shown in Figure 6C, immunoblot analysis of cell lysates revealed that CCCP treatment notably decreased the polyubiquitination of PINK1 in the presence of USP20-WT upon CCCP treatment (Figure 6C), contributing mainly to the accumulation of PINK1 levels. The overall data indicate that USP20 facilitates the deubiquitination of PINK1 after treatment with CCCP.

**Figure 6.**
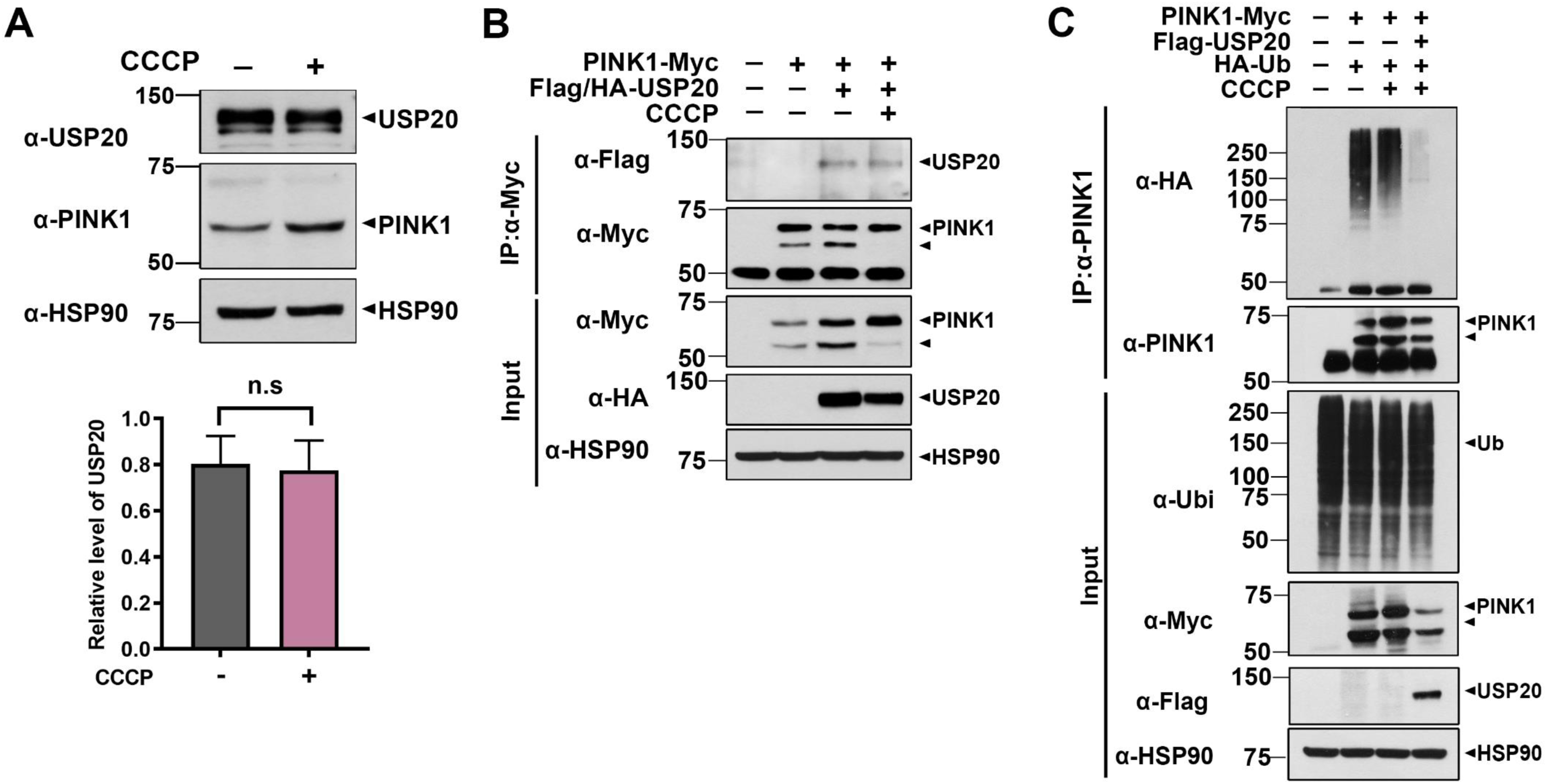
USP20 catalyzes deubiquitination of PINK1 upon CCCP treatment. **(A)** Where specified, HEK293 cells were treated for 4 h with 10 μM CCCP, and cell lysates were immunoblotted with anti-USP20 antiserum. Relative USP20 levels were quantified and are presented as the mean ± S.D. of three independent experiments (N.S., not significant). **(B)** HEK293 cells were transfected for 24 h with plasmids encoding PINK1-Myc alone or in combination with Flag/HA-USP20, and treated 4 h with 10 μM CCCP. Cell lysates were immunoprecipitated using anti-Myc antibody, followed by immunoblotting with the indicated antibodies. **(C)** HEK293 cells were transfected for 24 h with plasmids encoding Myc-PINK1 alone or in combination with Flag-USP20, and treated for an additional 6 h with 10 μM MG132. Cells were then treated 4 h with 10 μM CCCP. Cell lysates were immunoprecipitated using an anti-PINK1 antibody, followed by immunoblotting with the indicated antibodies. HSP90 served as a loading control.

### USP20 facilitates CCCP-induced mitophagy by promoting PINK1 accumulation

Building upon our prior investigation into the role of USP20 in elevating PINK1 levels and promoting autophagy induction, we conducted further analyses to explore whether USP20 induces mitophagy and contributes to the restoration of MMP. To assess MMP, we employed the commonly used JC-1 dye, which exhibits green fluorescence under CCCP treatment, indicating membrane depolarization and red fluorescence under normal condition. In this assay, HEK293 cells were transfected with USP20 and/or treated with CCCP. Staining with JC-1 dye revealed that the presence of USP20 significantly rescued the CCCP-induced loss of MMP (Figure 7A). Furthermore, we compared its effects in wild-type PINK1-expressing and *PINK1*-null MEF cells. The results demonstrated that overexpression of USP20 had a restorative effect on CCCP-induced MMP loss in wild-type PINK1-expressing MEF cells, while no significant difference was observed in *PINK1*-null MEF cells (Figure 7B).

**Figure 7.**
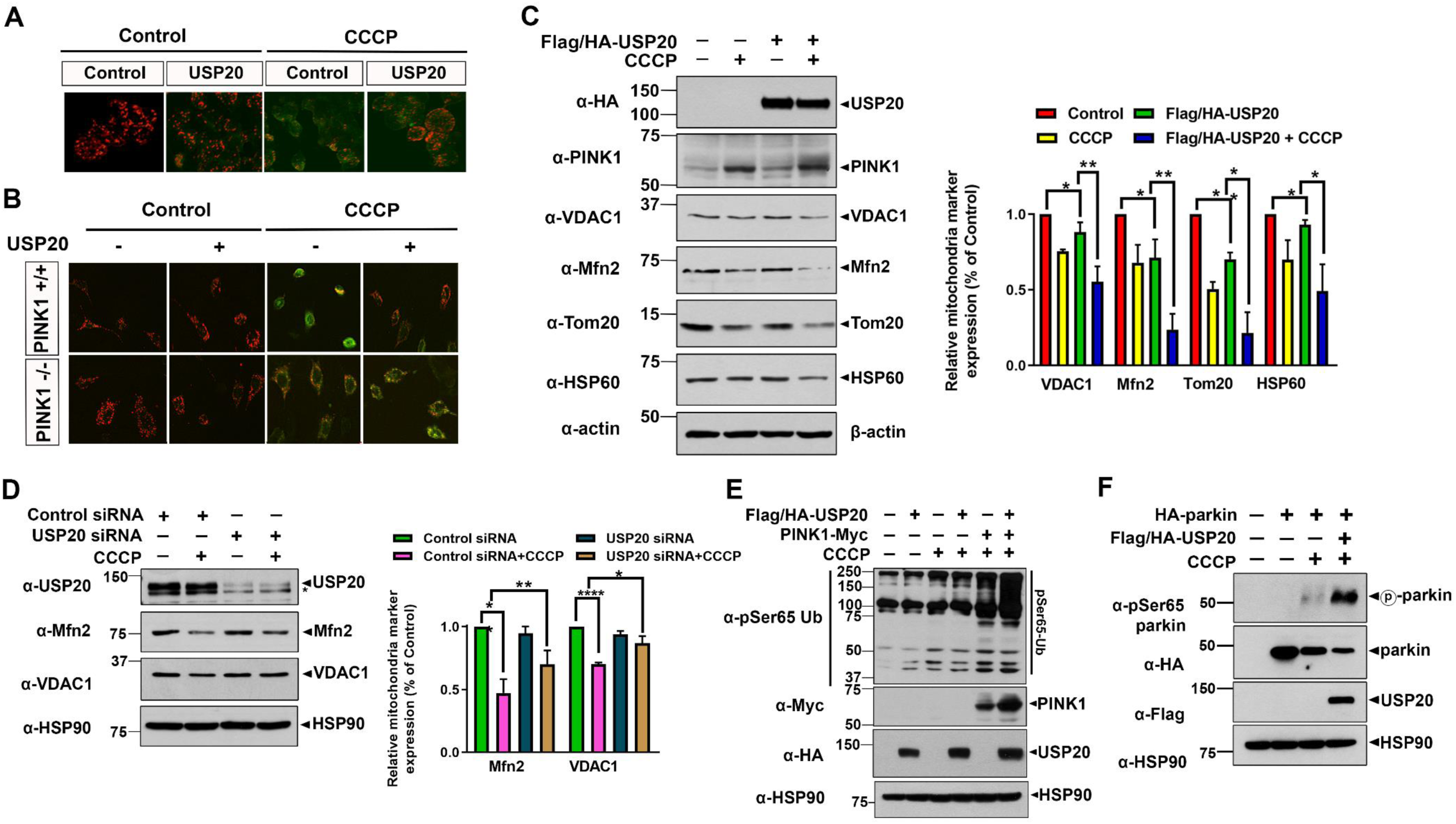
USP20 promotes CCCP-induced mitophagy. **(A)** HEK293 cells were mock-transfected or transfected for 24 h with a plasmid encoding Flag/HA-USP20. The cells were then treated 4 h with 10 μM CCCP and stained with the JC-1 fluorescent dye. **(B)** Where indicated, PINK1^+/+^ or PINK1^-/-^ MEFs were mock-transfected or transfected for 24 h with a plasmid encoding Flag/HA-USP20. Cells were then treated 4 h with 10 μM CCCP, and stained with the JC-1 dye. The stained cells were analyzed by confocal microscopy. **(C)** HEK293 cells were transfected for 24 h with a plasmid encoding Flag/HA-USP20, and treated with 10 μM CCCP for 4 h. Cell lysates were immunoblotted with the indicated antibodies. Each level of mitochondrial markers was presented as the mean ± S.D. of three independent experiments (\*\**p* ≤ 0.001; \**p* ≤ 0.05). **(D)** HEK293 cells were transfected for 48 h with control siRNA (CTL) or USP20-siRNA, and treated 4 h with 10 μM CCCP. Cell lysates were immunoblotted and with the indicated antibodies. Data are presented as the mean ± S.D. of three independent experiments (\*\*\**p* ≤ 0.0001; \*\**p* ≤ 0.001). **(E)** HEK293 cells were transfected for 24 h with plasmids encoding PINK1-Myc alone or in combination with Flag/HA-USP20, and treated 4 h with 10 μM CCCP. Cell lysates were immunoblotted with the indicated antibodies. **(F)** HEK293 cells were transfected for 24 h with plasmids encoding HA-parkin alone or in combination with Flag/HA-USP20, and treated for additional 6 h with 10 μM CCCP. Cell lysates were immunoblotted with the indicated antibodies. Actin and HSP90 served as loading controls.

Additionally, we investigated the impact of USP20 on mitochondrial protein levels under CCCP treatment. As anticipated, several mitochondrial membrane protein levels (i.e., VDAC1, Mfn2, Tom20, HSP60) exhibited a significant decrease in the presence of USP20 with CCCP treatment (Figure 7C). In contrast, cells with siRNA-mediated USP20 knockdown displayed a slight increase in these protein levels compared to cells transfected with control siRNA (Figure 7D). Given that the consecutive phosphorylation of ubiquitin and parkin at serine 65 (S65) residue is essential for the initiating mitophagy (Xu et al., 2021), leading to the translocation and activation of parkin, we further examined the effect of USP20 on the phosphorylation status of ubiquitin at S65 after CCCP treatment. Immunoblotting of cell lysates revealed that the phopho-S65 level of ubiquitin was substantially elevated when USP20 was co-transfected, and it was further augmented with CCCP treatment (Figure 7E). Subsequently, we examined the phosphorylation of parkin in HEK293 cells and found that USP20 significantly increased the phosphorylation of parkin at S65, compared to treatment with CCCP alone. (Figure 7F).

Collectively, these findings suggest that USP20 functions as a novel regulator, promoting PINK1 accumulation and stimulating mitophagy in response to CCCP treatment.

### Elevation of Familial PD-Linked PINK1 Mutant and Wild-Type PINK1 Levels by USP20

Finally, we explored the impact of USP20 on two familial PD-linked pathogenic mutants of PINK1, namely PINK1-G309D and PINK1-L347P, with substitutions at G309 and L347 by Asp and Pro, respectively (Figure 8A). Previous studies indicated that these two familial point mutants of PINK1 exhibit significantly reduced kinase activity and protein stability compared to wild-type PINK1 (Beilina et al., 2005). As depicted in Figure S4B, HEK293 cells were transfected with a plasmid encoding USP20-WT, PINK1-WT, PINK1-G309D, or PINK1-L347P alone or in combination. Immunoblot analysis of cell lysates revealed that the overexpression of USP20 increased the levels of both familial PINK1 mutants, and their enhanced levels were comparable to that of wild-type PINK1 (Figure 8B). These findings suggest that USP20 induces the accumulation of the two PD-associated PINK1 mutants alongside PINK1-WT.

**Figure 8.**
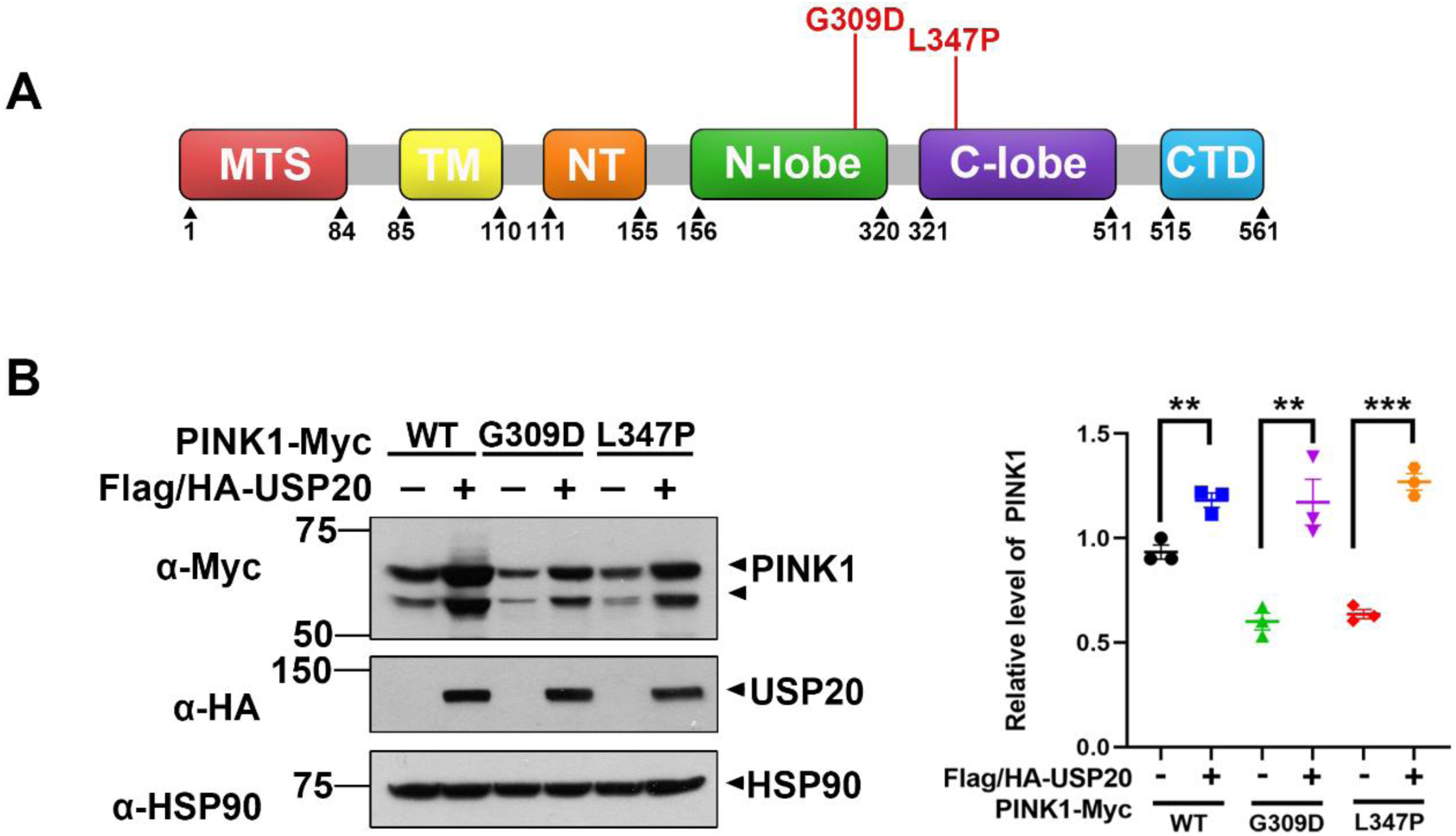
USP20 increases two familial PD-associated PINK1 mutants as well as its wild type form. **(A)** Schematic of two PD-linked mutations in PINK1. **(B)** Where specified, HEK293 cells were transfected for 24 h with plasmids encoding Flag/HA-USP20, wild-type PINK1-Myc, PINK1-Myc-G309D, or PINK1-Myc-L347P mutant alone or in combination. Cell lysates were immunoblotted with the indicated antibodies. Relative levels of PINK1 were quantified and the results are represented as the mean ± S.D. of three independent experiments (\*\*\**p* ≤ 0.0001; \*\**p* ≤ 0.001). HSP90 served as a loading control

Overall, our data suggest that USP20 may represent a novel therapeutic target for PD by facilitating the deubiquitination of PINK1 and stimulating its downstream mitophagic flux activity, thereby influencing the potential role of USP20 in PD pathogenesis.

## Discussion

The deubiquitinating enzymes play a critical role in ubiquitin signaling processes, having the function of removing ubiquitin from substrates (Estavoyer et al., 2022). The significance of DUBs has been highlighted not only in the pathogenesis of neurodegenerative diseases (NDDs) but also in cancer. In the context of NDDs, many studies have revealed that AD is closely associated with the accumulation of β-amyloid peptide (Aβ) and the microtubule-binding protein tau in the brain (Murphy and LeVine, 2010). The amyloid precursor protein (APP) binds to USP25 and becomes degraded via the UPS pathway under ER stress (Jung et al., 2015). In addition, the inhibition of USP13 increases proteasome activity and decreases hyper-phosphorylated tau (Liu et al., 2019). Overall, the deubiquitination of several substrates targeted by numerous DUBs has been shown to influence NDDs. These studies further demonstrate them to be promising candidates for inhibiting the progression of NDDs, including PD. The current study could provide additional evidence, supporting the idea that another DUB, USP20, is critical for modulating PINK1-mediated neuronal cell death or the progression of PD.

Since mitochondrial dysfunction is a common primary or secondary contributor to various NDDs, including PD, it is crucial to comprehend the precise mechanism of the mitophagy pathway and its associated operators/regulators for a comprehensive understanding of PD pathology. To date, two key players, PINK1 and parkin, are recognized for their vital roles in executing and regulating mitophagy by selectively eliminating damaged mitochondria (Palikaras and Tavernarakis, 2014).

Given the functional significance of mitophagy in PD pathology and its link to DUBs, multiple studies have revealed the direct or indirect interaction between parkin and several DUBs. For example, USP8 acts as a direct and positive regulator for parkin, selectively increasing its protein levels and eliminating the K6-linked ubiquitin chain from parkin (Durcan et al., 2014; Tsefou and Ketteler, 2022). While the exact working mechanism remains unclear, the identified indirect link between parkin and a few USPs influences the functional properties of parkin, such as mitochondrial recruitment, rather than the accumulation of parkin levels, ultimately promoting mitophagy. For example, the depletion of *USP30* has accelerated the recruitment of parkin into mitochondria, leading to the removal of K6-linked ubiquitin chains and promoting mitophagy (Park et al., 2021; Wang et al., 2015; Luo et al., 2021). Additionally, USP33 directly interacts with parkin, and the knockdown of *USP33* increases K63-linked ubiquitination of parkin under CCCP treatment, thereby enhancing mitophagy (Niu et al., 2022). The therapeutic potential of targeting DUBs lies in their ability to influence mitophagy. By inhibiting specific DUBs, researchers aim to enhance the degradation of damaged mitochondria, thereby alleviating the cellular stress associated with PD (Park et al., 2021).

Despite the identification of multiple positive deubiquitinating enzymes for parkin, the positive regulatory mechanism for PINK1 stability remains unknown. While a few Ub E3 ligases that mediate the proteolytic degradation of PINK1 have been identified, its positive regulator(s) still remain elusive. Hence, there is a necessity to explore PINK1-targeting DUBs, as PINK1 serves as an upstream factor of parkin. Here we discover, for the first time, that the deubiquitinating enzyme USP20 promotes the accumulation of PINK1 in mammalian cells, providing insight into the positive regulatory mechanism and influencing the expression of PINK1 in various cellular contexts. Furthermore, additional *in vitro* data support the notion that USP20 has the potential to modulate mitochondrial quality control by regulating PINK1 and PINK1-mediated mitophagy, thereby contributing to our understanding of the pathogenesis of PD.

In addition to PINK1, many recent studies suggested that several PD-related genes, such as α-synuclein, FBXO7, and LRRK2, are also modulated by the actions of DUBs. For example, the abnormal aggregation of α-synuclein, the major constitute of Lewy body, is known to be strongly correlated with PD pathogenesis (Gómez-Benito et al., 2020). The knockdown of *USP9X* increased the amount of aggregated α-synuclein, thereby exacerbating the pathogenesis of PD (Rott et al., 2011). Moreover, *USP13*-knockdown increases α-synuclein ubiquitination in a parkin-independent manner, regulating α-synuclein-induced neuronal death (Liu et al., 2019). Additionally, mutations in FBXO7 gene lead to autosomal recessive PD, and FBXO7 has been demonstrated to facilitate mitochondrial homeostasis and mitophagy, which is antagonized by the action of USP30 upon a yet unclear stimulus (Sanchez-Martinez et al., 2023). Moreover, we have recently reported that USP7 directly interacts with FBXO7, positively regulating the stability of FBXO7 through K48-linked deubiquitination. (Lee and Chung, 2023). The overall studies verified a strong association between USPs and several PD-related genes, highlighting the crucial role of DUBs in regulating various cellular activities. Based on these results, the present study further demonstrates the correlation between PINK1, another key-contributing member of familial PD cases, and USP20, a member of DUB family, stabilizing the half-life of PINK1 and consequently positively stimulating mitophagy. The discovery of USP20 as PINK1 modulator, confirms the presence of complete and reversible regulatory mode of PINK1 stability and its novel player, providing an indirect evidence for functional importance of PINK1 in mitophagy and the pathology of PD.

Regarding the functional importance of USP20, it plays a key role in regulating various cellular pathways and cell viability. Specifically, USP20 deubiquitinates β-catenin, influencing its protein stability and promoting cell proliferation, thus establishing a critical regulatory mechanism in tumorigenesis (Wu et al., 2018). Additionally, USP20 has been shown to regulate cancer metastasis. For instance, the knockdown of USP20 inhibits cell invasion by deubiquitinating SNAI2, potentially suppressing cancer metastasis (Beilina et al., 2005). Moreover, a recent discovery indicates that USP20 has a close relationship with autophagy-related proteins, such as ULK1 and p62. ULK1 plays a role in forming autophagosomes, which ultimately fuse with lysosomes and mediate the degradation of unnecessary products. Similarly, USP20 interacts with ULK1 at basal levels, controlling the initiation of autophagy by maintaining the basal expression level of ULK1 (Kim et al., 2018). Furthermore, USP20 binds with p62/sequestosome-1 and regulates its stability through deubiquitinating K48 residues, delivering ubiquitinated cargo to lysosomes and undergoing autophagic degradation (Ha et al., 2020; Lim et al., 2020). Therefore, the deubiquitinating activities of many DUBs play a critical role in maintaining cellular homeostasis and viability in cancer and autophagy. While no direct relationship between DUBs and PD has been established previously, this study identifies USP20 as a novel DUB target for PD.

In conclusion, this study establishes that USP20 plays a crucial role in catalyzing the deubiquitination of PINK1, preventing its degradation via UPS and functioning as a positive mediator for PINK1. Furthermore, our findings demonstrate that the accumulation of PINK1 facilitated by USP20 significantly enhances mitophagy, thereby influencing mitochondrial dynamics. Given the neuroprotective function of PINK1 and the various forms of its loss-of-function mutations prevalent in most familial PD cases, the insights gained from this study regarding the novel action of USP20 could pave the way for innovative approaches in PD therapeutics.

## Data availability

The data used and analyzed during the current study are available from the corresponding author on reasonable requests.

## Acknowledgements

We thank J. Chung, J. Song, M.R. Cookson, and S.H. Park for providing plasmids, and J. Shen for providing MEFs derived from *PINK1*-null mice, respectively.

## Author contributions

Ga Hyun Park performed the experiments, including DNA transfection, western blotting, *in vitro* GST-pull down, and PLA, and analyzed the data. She also wrote the manuscript. Hye In Park conducted western blotting and measured mitochondrial membrane potential, and analyzed the data. Donghyuk Shin conducted alphafold prediction and analyzed the data. Kwang Chul Chung participated in the conception and design the experiments, as well as their analysis and interpretation. He also wrote the article, obtained funding, and carries overall responsibility for the data. All authors have read and approved of this manuscript.

## Supplementary material

This article contains Supplementary material.

