## Supplementary Data for "Ubiquitin-specific protease 20 promotes CCCP-induced mitophagy through deubiquitination and stabilization of serine/threonine protein kinase PINK1"

### Contents

1. **Figure S1.** USP20 increases the protein level of PINK1 in mammalian cells.
2. **Figure S2.** USP20 increases the protein stability of PINK1.

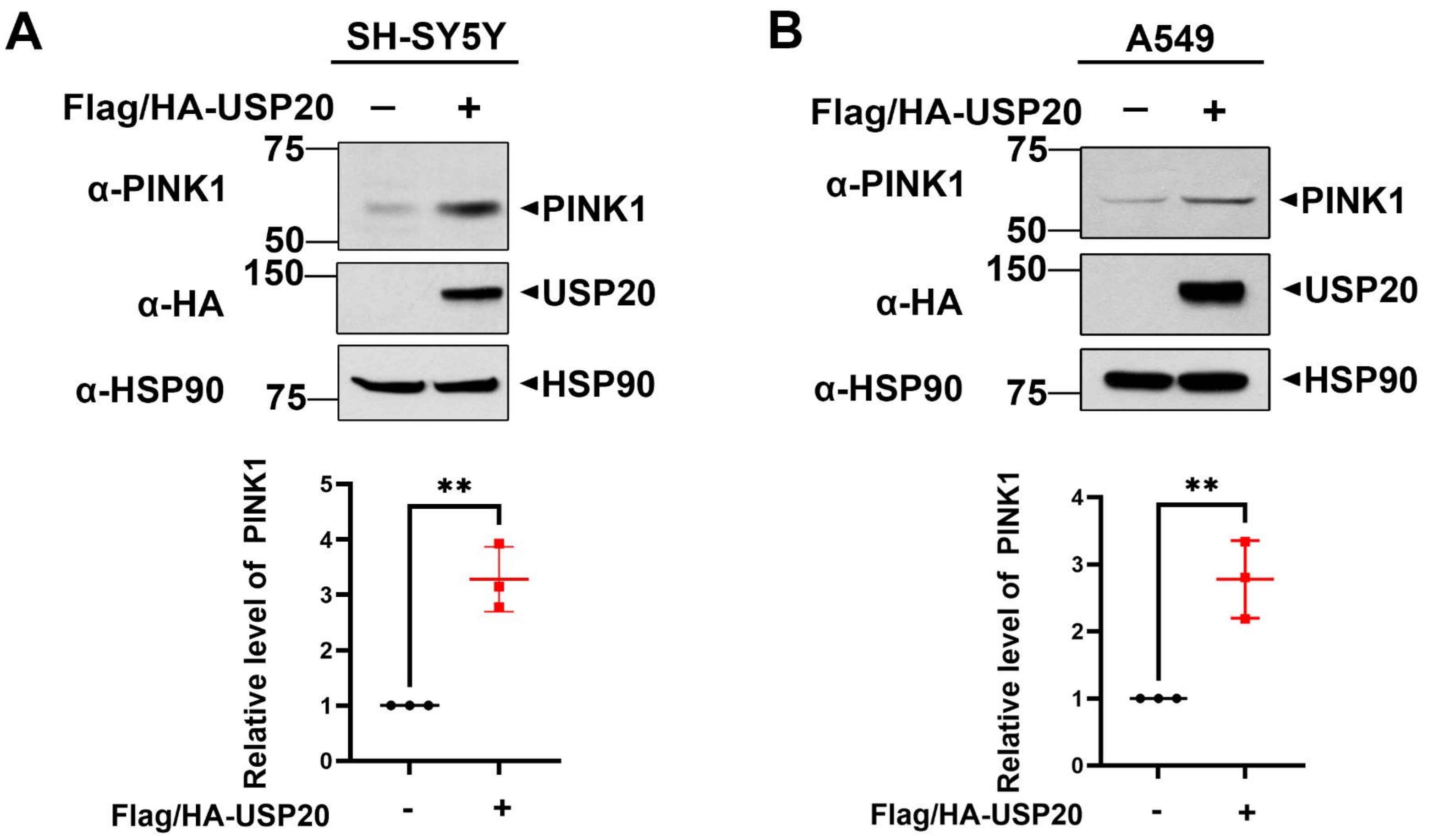


**Figure S1. USP20 increases the protein level of PINK1 in mammalian cells.** (A) SH-SY5Y cells were transfected for 24 h with plasmids encoding Flag/HA-USP20-WT, and cell lysates were immunoblotted with anti-PINK1 antibodies. Relative PINK1 levels were quantified and the results are presented as the mean ± SD of three independent experiments (**$p \leq 0.001$). (B) A549 cells were transfected for 24 h with plasmids encoding Flag/HA-USP20-WT, and cell lysates were immunoblotted with anti-PINK1 antibodies. Relative PINK1 levels were quantified and the results are presented as the mean ± SD of three independent experiments (**$p \leq 0.001$).

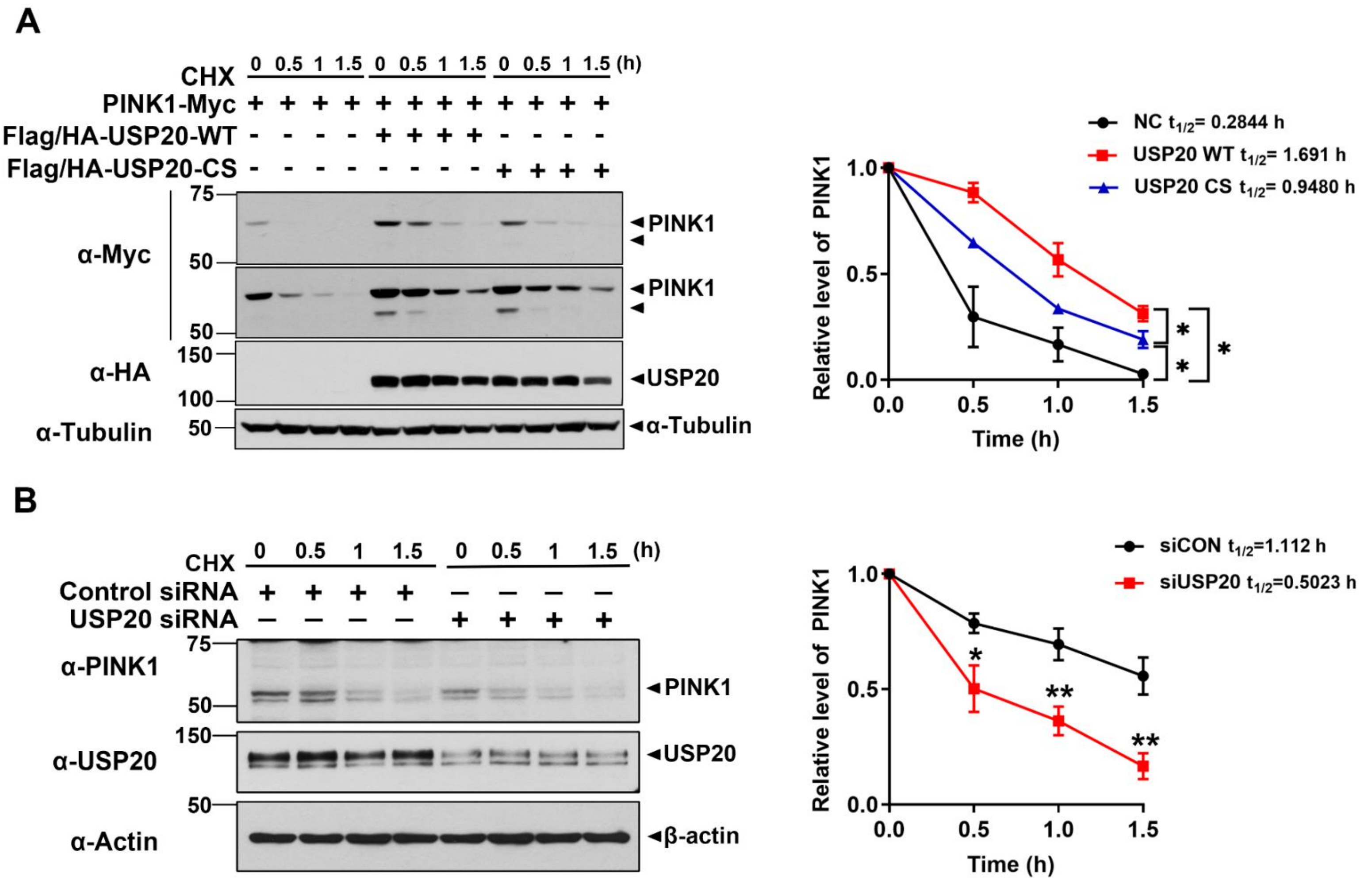


**Figure S2. USP20 increases the protein stability of PINK1.** (A) HEK293 cells were transfected for 24 h with plasmids encoding wild type PINK1-Myc, Flag/HA-USP20-WT, or Flag/HA-USP20-CS alone or in combination. Cells were treated for the indicated times with 100 μg/ml of cycloheximide, and cell lysates were immunoblotted with the indicated antibodies. Relative PINK1 levels were quantified, and the results are presented as the mean ± S.D. of three independent experiments (*$p \leq 0.05$). (B) HEK293 cells were transfected for 48 h with control siRNA or *USP20*-siRNA. Cells were treated for the indicated times with 100 μg/ml cycloheximide, and cell lysates were immunoblotted with the indicated antibodies. Relative levels of PINK1 were quantified and the results are presented as the mean ± S.D. of three independent experiments (**$p \leq 0.001$; *$p \leq 0.05$). Tubulin and actin served as loading controls.
